# *Prototaxites* was an extinct lineage of multicellular terrestrial eukaryotes

**DOI:** 10.1101/2025.03.14.643340

**Authors:** Corentin C. Loron, Laura M. Cooper, Sean McMahon, Seán F. Jordan, Andrei V. Gromov, Matthew Humpage, Laetitia Pichevin, Hendrik Vondracek, Ruaridh Alexander, Edwin Rodriguez Dzul, Alexander T. Brasier, Alexander J. Hetherington

## Abstract

*Prototaxites* was the first giant organism to live on the terrestrial surface, reaching sizes of 8 metres in the Early Devonian. However, its taxonomic assignment has been debated for over 165 years^1–7^. Tentative assignments to groups of multicellular algae or land plants^1,2,8–11^ have been repeatedly ruled out based on anatomy and chemistry^5,7,11–16^ resulting in two major alternatives: *Prototaxites* was either a fungus^5,6,17–22^ or a now entirely extinct lineage ^7,16,23^. Recent studies have converged on a fungal affinity^5–7,17–20,22^. Here we test this by contrasting the anatomy and molecular composition of *Prototaxites* with contemporary fungi from the 407-million-year-old Rhynie chert. We report that *Prototaxites taiti* was the largest organism in the Rhynie ecosystem and its anatomy was fundamentally distinct from all known extant or extinct fungi. Furthermore, our molecular composition analysis indicates that cell walls of *P. taiti* include aliphatic, aromatic, and phenolic components most similar to fossilisation products of lignin, but no fossilisation products characteristic of chitin or chitosan, which are diagnostic of all groups of extant and extinct fungi, including those preserved in the Rhynie chert. We therefore conclude that *Prototaxites* was not a fungus, and instead propose it is best assigned to a now entirely extinct terrestrial lineage.

---

*Prototaxites* as previously identified in the Rhynie chert constitutes one species, *P. taiti*. The original description of *P. taiti*^24^ was based on a small fragment preserved in two consecutive thin sections, which consists of a mass of wide and narrow tubes and two darker roughly spherical regions termed medullary spots (**Figs. 1a, b**). A second fragment of *P. taiti* was identified in two thin sections cut from the same block and interpreted as a peripheral region (**Fig. 1c**). The peripheral region was re-investigated in 2018 and proposed to be a fertile structure composed of tubes resembling asci with small ascospores^6^. This description led the authors to assign *P. taiti* to the Ascomycota, as a “basal member” of this extant fungal group^5–7,17–20,22^. This assignment remains contentious as *P. taiti* possesses a unique combination of characteristics^7^ and most critically the proposed fertile fragment lacks organic connection to the vegetative material^23^. Based on our re-investigation of the *P. taiti* type material we recognise three key points. First, until connection can be demonstrated between the proposed fertile and the vegetative material, we agree with Edwards^25^ that they should be interpreted separately. Second, the vegetative type material is tiny when compared to the tree-sized *Prototaxites* known elsewhere from the Devonian^1–5,10,18,20^. Establishing if *P. taiti* was tiny or if this small size is the result of preservational bias is essential for interpreting its ecology. Finally, we recognise that the Rhynie chert offers a unique opportunity to test the taxonomic assignment of *Prototaxites* by contrasting its chemistry with a diversity of prokaryotic and eukaryotic fossils that have experienced the same diagenetic history. Based on these points we conducted an extensive re-examination of *P. taiti*, leading us to reject the most widely held hypothesis that *Prototaxites* was a fungus^5–7,17–20,22^.

**Fig 1.**
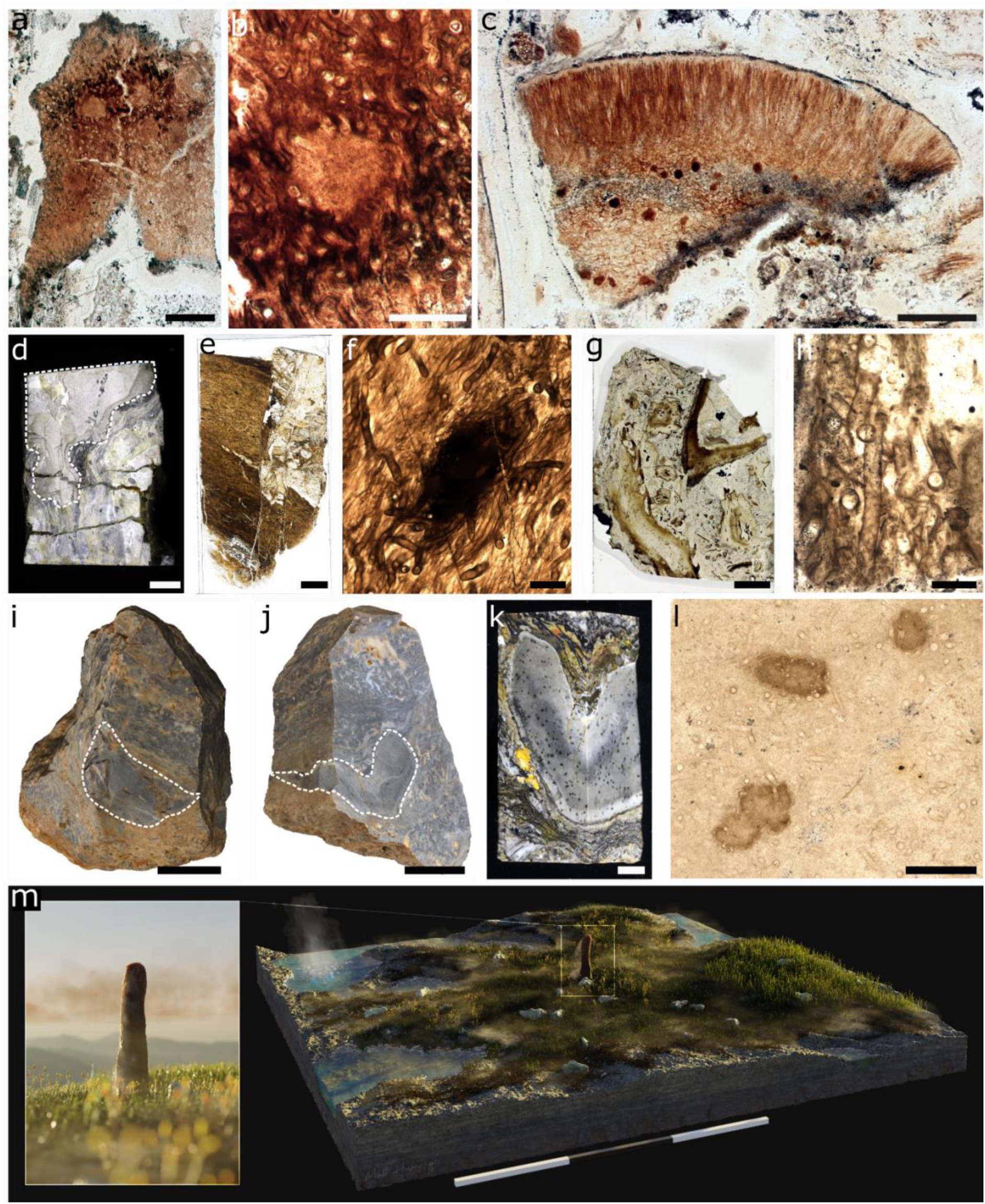
| *Prototaxites taiti* material from the Rhynie chert. **a-c**, Images of two of the four thin sections containing the fragments that constitute the *P. taiti* type material, including medullary spots (**b**) and peripheral region (**c**). **d-l:** *P. taiti* material used in this study. **d,e: d**, Lyon 156 with *P. taiti* highlighted in dashed lines. **e**, thin section produced from the block in (**d**) showing the fractured *P. taiti* specimen. **f**, Magnified image of the thin section in (**e**) showing the characteristic tubes and medullary spots of *P. taiti*. **g, h: g**, thin section made from Lyon 48 with *P. taiti* in dashed box. **h**, detail of thin section in (**g)** showing the tubes. **i-m**: Imaging and reconstruction of a large, exceptionally well-preserved *P. taiti* from NSC.36. **i**, Photogrammetry model of NSC.36 before cutting with surface exposed *P. taiti* circled by dashed line. **j**, Photogrammetry model of NSC.36 after initial cutting of the block with *P. taiti* circled by a dashed line. **k**, Block of NSC.36 from which thin sections were produced, showing medullary spots throughout the body. **l**, thin section taken from the block in **(k)** showing characteristic tubes and medullary spots of *P. taiti.* **m,** Artist reconstruction of *P. taiti* within the Rhynie ecosystem including hypothesised reconstruction of the aerial portion. Illustration by Matthew Humpage, Northern Rouge Studios. Scale bars: 3m (**m**), 3 cm (**i**), 2cm (**j**), 1cm (**d**), 5mm (**e, g, k**), 1mm (**c**), 500µm (**a, l**), 200µm (**b, f**), 100µm (**h**). Specimen accession codes: GLAHM Kid 2523 (**a**, **b**), GLAHM Kid 2525 (**c**), Lyon 156 (**d**), Lyon 156 MPEG0078 (**e, f**), NSC.36 (**i-k**), NMS G.2024.5.7 (**l**).

Our investigation of *P. taiti* draws upon three specimens: a large but highly fractured specimen in block Lyon 156 described in Edwards^25^ (**Fig. 1d-f**), a small fragment of *P. taiti* identified in block Lyon 48 (**Fig. 1g, h**), and a large and exceptionally well-preserved specimen identified in block NSC.36 (**Fig. 1 i-l**). Owing to the large size and exceptional preservation of the *P. taiti* specimen in NSC.36, hence referred to as NSC.36, we focus our description on this specimen. NSC.36 is roughly cylindrical, 5.6 cm at its widest, and extends obliquely through the entirety of the block (6.9 cm). To our knowledge this is the largest *P. taiti* specimen reported from the Rhynie chert^6,24,25^ and as the specimen is incomplete, we infer that *P. taiti* was the largest known organism in the Rhynie ecosystem. The block was initially cut, revealing that NSC.36 is a boomerang shape in transverse section (**Fig. 1j**), the longer arm being 3.9 cm long, the other 3.6 cm. Orientation of the specimen was determined using geopetal features (**Extended Data Fig. 1; Supplementary Material**).

Internally, NSC.36 comprises a light brown body with abundant and evenly distributed, dark brown, roughly spherical medullary spots^3,4,6,24,25^, 200 to 600 µm in diameter (**Fig. 1 k,l**), and a black carbonised layer delineates the specimen from the surrounding substrate (**Fig. 1k**). After describing the macro-scale features, thin sections were produced to investigate the anatomy.

At the micro scale, NSC.36 is composed of tubes with different anatomies (**Fig. 2a**). We identified three distinct tube types in the main body. Type 1 tubes are small in diameter (∼10 µm), thin (<1 µm) and smooth-walled, septate with pores, sinuous, occasionally branching, and tend to be oriented with the long axis of the specimen (**Fig. 2b**). Type 1 tubes constitute ∼75% of the body volume. Type 2 tubes are larger in diameter (ranging from 20–40 µm), have smooth, double layered (∼2 µm thick) walls, and are aseptate, sinuous and unbranched (**Fig. 2c**). Type 2 tubes are similarly oriented with the long axis of the specimen and are intermingled with the type 1 tubes. Type 2 tubes constitute ∼20% of the body volume. Type 3 tubes are the largest in diameter (∼40 µm), thick-walled (∼2 µm) with weak annular thickenings, and are aseptate and unbranched (**Fig. 2d**). Type 3 tubes also align with the long axis and comprise ∼5% of the body by volume (**Fig. 1d, h**).

**Fig. 2.**
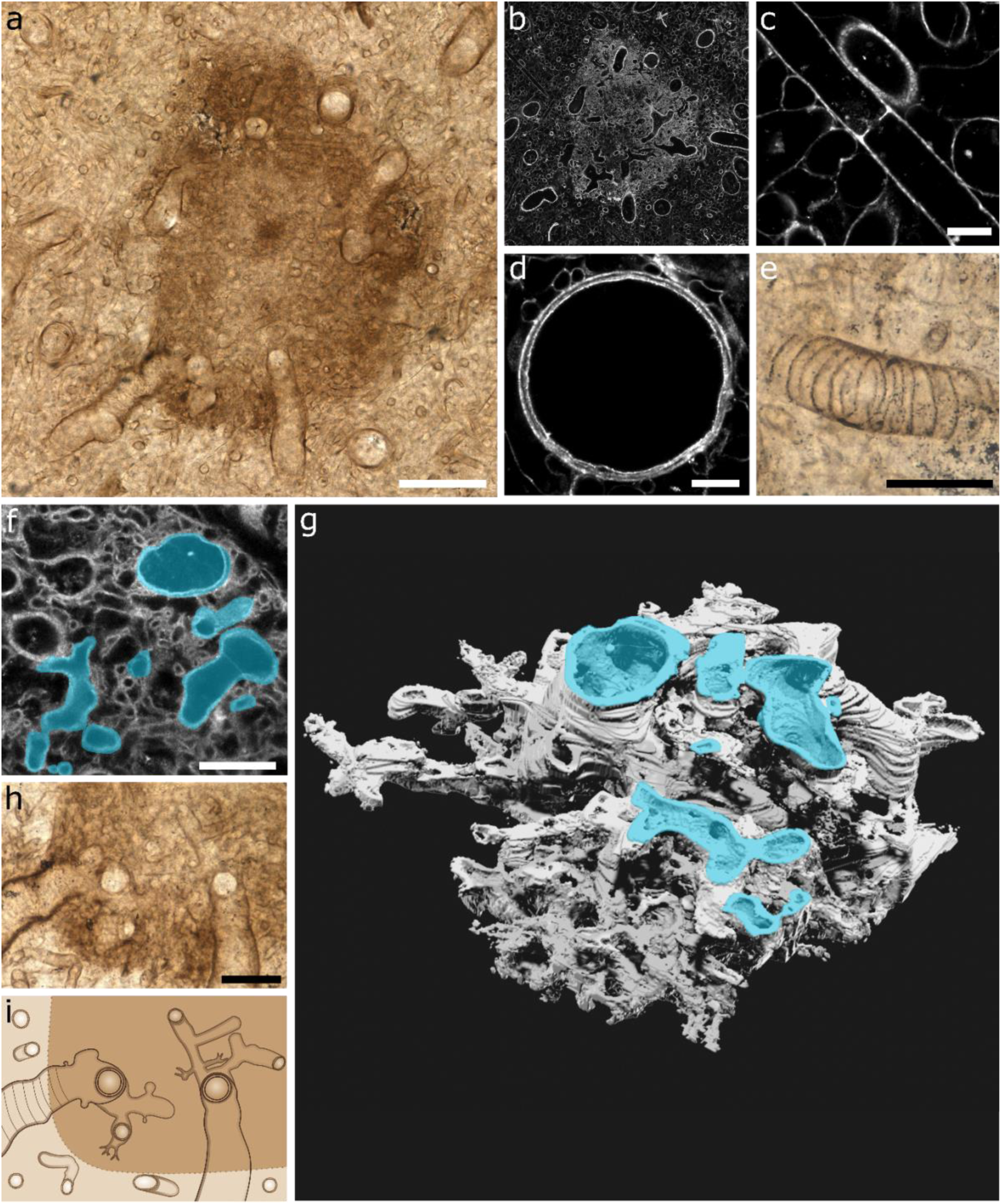
| The medullary spots and tube types of *Prototaxites taiti* are morphologically distinct from anything observed in extinct or extant fungal groups, and do not support a fungal affinity for *Prototaxites*. **a,** Transmitted light image showing a medullary spot within the body of *P. taiti*. **b,** The same medullary spot imaged using CLSM, showing the spot to be composed of densely packed fine tubes contrasting with the less densely packed body. **c-e,** Details of tubes types 1-3 seen in the body of *P. taiti*: small diameter type 1 tube with septal pore **(c)**, larger diameter type 2 tube **(d),** and type 3 tube with annular thickenings **(e). f-h,** Airyscan CLSM three-dimensional imaging reveals that at the medullary spot region all tube types are connected through a highly branched network. Tubes of a variety of morphologies (highlighted in cyan in **f** and **g**) were found to be connected to each other in a dense and fine branching network through the construction of a 3D SPIERS model (**g**) using Airyscan CLSM z-stack data (the first image in the stack is shown in **f**). Examination of the spot region (**h**) supports the interconnection of all tube types through fine branching at the medullaryspots, as shown in the schematic in (**i**). Scale bars: 100µm **(a)**, 50µm **(e, h**), 20 µm **(f**), 10µm **(c,d)**. Specimen accession code: NMS G.2024.5.7.

The identification of three tube types does follow previous anatomical descriptions of other *Prototaxites* species, and this observation was taken by Hueber^5^ to support the hypothesis that *Prototaxites* was a basidiomycete. In this group, fruiting bodies can have a trimitic hyphal construction – composed of binding, generative and skeletal hyphae^26, 27^. The type 1 and 2 tubes described here do fit with Hueber’s description of binding and generative hyphae. However, our description of type 3 tubes, particularly the annular thickenings (also observed in Lyon 156 (**Extended Data Fig. 2)** and other nematophytes^28^), is not compatible with the skeletal hyphae of extant fungi, which lack annular thickenings^29^. Furthermore, in extant fungi, annular thickenings are only found in elaters^30^ rather than constituting the internal body. Therefore, the anatomy of type 3 tubes does not support assignment to any modern fungal group.

*P. taiti* also differs from extant fungi due to the occurrence, anatomy, and development of the medullary spots. Optical and Airyscan Confocal Laser Scanning Microscopy (CLSM) examination of the medullary spots of NSC.36 revealed these structures to consist of a dense network of the type 1-3 tubes described above, as well as very fine (as small as 1 µm in diameter) tubes (**Fig. 2e)**. With the exception of the fine branching, the tube types in spots are anatomically identical to those found in the body, but to investigate if they were molecularly identical, we obtained high resolution maps with synchrotron Fourier Transform Infrared Spectroscopy (FTIR) of both the body and the medullary spots. Both regions were similar across the full length of their available spectra (3000-1400 cm-1) (**Supplementary** Figs 3-5), consistent with a similar molecular composition. Having demonstrated that the body and medullary spots were anatomically and chemically similar, we aimed to define the connection between tube types within medullary spots using CLSM. Airyscan CLSM z-stack data was used to produce a 3D reconstruction of the tube network in a single medullary spot and determine the connections between tubes (**Fig. 2f**). Using our 3D model, we found connections between all three tube types in the medullary spots, and demonstrated their highly interconnected nature and therefore complex development (**Fig. 2g-i)**. This finding is in marked contrast to extant fungi for two reasons. Firstly, there are no structures analogous to medullary spots in extant fungi. Hueber^5^ proposed similarities between medullary spots and the coltricoid hyphae of some basidiomycetes, however there are significant differences. Namely, in coltricoid hyphae, fine branching is exclusively produced from lateral emergences of wide hyphae that are aligned together^29^, and branching order is highly specific, with only generative hyphae giving rise to other hyphal types (binding and skeletal^27^). This is contrasted with medullary spots, where fine branching does not predominately occur from lateral emergences of larger tubes, there is no directional alignment of tubes, and no branching order can be discerned (**Fig. 2h, i**).

Our investigation of NSC.36 therefore demonstrates that *P. taiti* was the largest organism in the Rhynie ecosystem (**Fig. 1m**), and that it is anatomically inconsistent with all known extant and fossil fungi.

To further test the fungal affinity of *P. taiti*, we contrasted its molecular composition with that of contemporary fossil fungi from the Rhynie chert using Attenuated Reflectance FTIR (ATR-FTIR). That all samples have experienced the same diagenetic history enables us to make side-by-side comparisons of molecular composition between *P. taiti* and other taxa, while baselining for the influence of diagenesis. Twelve ATR-FTIR acquisition spots were measured across the three distinct *P. taiti* specimens. We hypothesised that if *P. taiti* was fungal, fossilisation products of amino-glucan sugars would be present in its cell walls, derived from an original chitin composition (a polymer of N-acetyl glucosamine), like that found in fungi and arthropods from the Rhynie chert^31^. To test this hypothesis, we combined the previously assembled dataset of 47 samples, representing six higher taxonomic groups^31^, with 55 additional samples from these six groups and *P. taiti* (See **Supplementary data** for full spectra for each sample, images of all sample spots, and details of taxonomic assignment). We then developed a novel analysis pipeline comprising two key steps: data exploration (Step 1) and modelling (Step 2) (**Extended Data Fig. 3; Extended Methods**). Step 1 aimed to test the hypothesis that biologically informative spectral bands correlated with taxonomic classification, and to select the most relevant spectral features for further modelling. In Step 2 we built classification models using supervised machine learning approaches to provide a robust statistical framework to accept or reject the taxonomic classification of each sample based on the selected spectral features.

We first performed a Canonical Correspondence Analysis (CCA), on the absorption bands selected using Principal Component Analysis (PCA) (see **Extended Methods**). The CCA (**Fig. 3**) shows a good separation between taxonomic groups in the ordination space and demonstrates clear correlations between spectral features and taxonomic assignment. Fungi, arthropods, peronosporomycetes (oomycetes) and amoeba are all strongly positively correlated with bands characteristic of the fossilisation products of sugar-protein compounds, especially the carbonyl and nitrogen moieties^32,33^, which would be expected from amino-glucan-rich precursors^31,34^ like chitin. By contrast, bacteria are positively correlated with aliphatic moieties. *P. taiti*, and to a lesser extent plants, are negatively correlated with aliphatic moieties. We conclude that there is a correlation between the fossilisation products of an organism and their taxonomic classification, as previously reported^31^, and suggest that the molecular fingerprint of *P. taiti* is distinct from that of the chitinous organisms in the Rhynie chert. Having established this correlation, we moved to step 2, where classification models were used to assign samples to a taxonomic group.

**Fig. 3.**
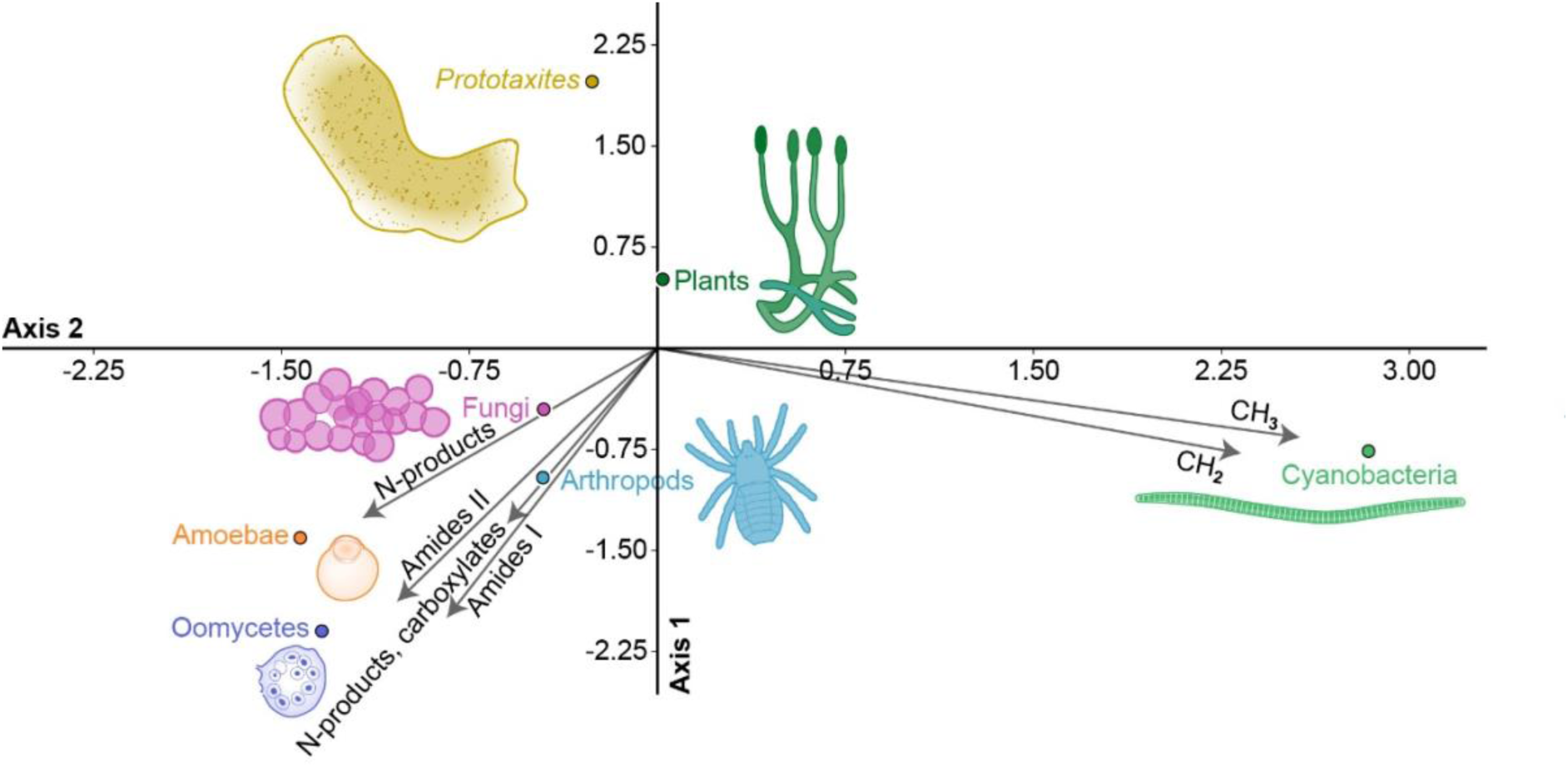
| The Canonical Correspondence Analysis (CCA) illustrates the correlation between informative spectral features and the fossils in the Rhynie chert. The CCA uses the most informative ATR-FTIR absorption bands obtained after dimension reduction and feature selection (bands 1, 2, 7, 10-12 in **Supplementary Table 1**, see **Methods, Extended Methods, Extended Data Figure 3**). It shows strong correlation between the fossilisation products of sugar-protein (Amide I, II, N-products and carboxylate) and Fungi, Arthropod, Peronosporomycetes (Oomycetes) and Amoebae. Bacteria show a strong correlation with aliphatic CH_x_ moieties. On the other hand, *Prototaxites taiti*, and land plants, shows a negative correlation with all these products. The CCA support the difference between the molecular composition of *P. taiti* and Fungi, and differences with the other fossils.

Unlike CCA, this approach leveraged the full spectra (reduced by PCA) and provided metrics for the confidence of assignment (see **Methods** and **Extended Methods**). We first trained our model on a dataset of *P. taiti* against fungi, then against all chitinous samples (fungi and arthropods). Support Vector Machine (SVM) models were produced for both datasets. All *P. taiti* samples were successfully identified in both test datasets. Globally, the models successfully discriminated *P. taiti*, with a discrimination accuracy of 91% against fungi and 93% against the chitinous group. All other performance metrics (recall, sensitivity, F1; see **Extended Methods**) scored above 91% and the Matthews Correlation Coefficient (MCC) — which measures the quality of the binary discrimination — was 0.85 for each dataset, respectively. These results confirm that the fossilisation products of *P. taiti*, as recorded in its molecular fingerprint, differ strongly from those of the chitinous organisms in the Rhynie chert. Having found no support for similarities between *P. taiti* and fungi, we then tested for similarities between *P. taiti* and bacteria, land plants, and a combined dataset of all other samples. In all cases, our models successfully discriminated *P. taiti* from all other groups (**Extended Methods**). Altogether, these results demonstrate that *P. taiti* cannot be assigned to any taxonomic group found in the Rhynie chert, and that the fossilisation products of chitin and chitosan, which are characteristic of extant and extinct fungi^35^, are absent from *P. taiti*.

These findings threw doubt upon the hypothesis that *P. taiti* was fungal and prompted a deeper molecular investigation of *P. taiti*. For the classification approach described above, thin section analysis was the only feasible approach to sample the full diversity of Rhynie chert taxa, some of which are microscopic. One potential drawback of this approach is that, as the samples are embedded in chert, the spectral bands produced by silica can mask some portions of the fossil spectra. This masking does not decrease the robustness of our approach, as its influence is identical across samples and silica bands are present in the features retained for the classification tasks (see **Extended Methods**). However, to characterise *P. taiti* in the absence of silica, we analysed the molecular composition of extracted fossil material. This was possible because block NSC.36 contained a *P. taiti* specimen large enough to permit extraction of organic content solely of *P. taiti* by acid maceration, and was surrounded by a peaty substrate containing predominantly plant, but also fungal and arthropod fossils (see **Fig. 1j, k; Extended Methods Fig. 1**). Although the small size and interbedded nature of these substrate fossils made a side-by-side comparison of *P. taiti* with these organisms intractable, we contrasted our extracted *P. taiti* sample with a bulk extraction of the substrate. The extracted *P. taiti* spectra (**Supplementary** Fig. 5) included bands characteristic of short/branched aliphatic moieties, carbonyl/carboxyl, aromatic and ether compounds, but not the ensemble of bands typically associated with polysaccharides^36^ (C-O-C, CC and CO stretching in the interval 1200 and 800 cm^-1^) and sugar-protein fossilisation products^31,34^. It is possible that polysaccharides and proteins were lost early in fossilisation and so did not form their characteristic C=O/N-rich fossilisation products^31^. However, our analysis reveals the presence of these products in the bulk extract of the peaty substrate. This indicates that these products are preservable in the Rhynie chert, suggesting an original absence of a strong polysaccharide contribution to *P. taiti* tissues. This further strengthening our argument that *P. taiti* was non-chitinous and accords with investigations of *Prototaxites* spp. from other localities^16^. Our new analyses together with the previous work of Abbott et al.^16^ allow us to conclude that *Prototaxites* had a conserved and distinct molecular fingerprint, and that it’s cell walls included a compound most similar to lignin, though distinct from the lignin of land plants (**Supplementary note**).

Finally, in order to further interrogate interpretation of *P. taiti* as an ascomycete^6^, we performed a biomarker analysis to test for the presence of perylene. Perylene is a known biomarker for fungi in the Rhynie chert^37^, and is derived from perylenequinones, pigment compounds produced predominantly by ascomycete fungi^38^. We contrasted samples consisting of purely of *P. taiti* with those of only the peaty substrate. Perylene was not detected in *P. taiti*, but was detected in the substrate (**Extended Data Fig. 4**), likely from the presence of ascomycetes (**Supplementary Material**). This biomarker analysis does not support an ascomycete affinity for *P. taiti*, and further demonstrates the distinctness of *P. taiti* from other organisms in the Rhynie chert.

We hypothesised that if *P. taiti* was a fungus, it would share molecular composition with contemporary fungi. All extant fungal clades – even the unresolved basal taxa (Rozellidae)^39^ – possess various amounts of chitin (or chitosan) during at least part of their life cycle, as well as beta-glucan^35,40^. The secondary loss and replacement of these foundational cell wall components would require significant alteration of major developmental pathways. Our data therefore do not support the placement of *Prototaxites* within Basidiomycota^5^ or Ascomycota^6^, even within a stem group^41^. For the same reason, it is also unlikely that *Prototaxites* is the result of an increase in complexity within an early-diverging fungal branch, e.g., Mucoromycota^22,42,43^. With no clear diagnostic ties to fungi, we conclude that *Prototaxites* was not a fungus.

Having found no support for the most widely held view that *Prototaxites* was fungal, we next reviewed possible placement in other higher taxonomic groups. No extant group was found to exhibit all the defining features of *Prototaxites*, namely: (1) formation of large multicellular structures of varied tube types; (2) a tube composition rich in aromatic-phenolic components; and, (3) a heterotrophic (likely saprotrophic) lifestyle (Table 1). Based on this investigation we are unable to assign *Prototaxites* to any extant lineage, reinforcing its uniqueness. We conclude that the morphology and molecular fingerprint of *P. taiti* is clearly distinct from that of the fungi and other organism preserved alongside it in the Rhynie chert, and we suggest that it is best considered a member of a previously undescribed, entirely extinct group of eukaryotes^7,8,16^.

**Table 1.** | No extant higher taxonomic group exhibits all the three defining features of *Prototaxites*.

| Candidate Taxonomic Group | Cell Anatomy and Structure | Main structural polymers | Carbon Assimilation Mechanism | Conclusion |
| --- | --- | --- | --- | --- |
| **Actinomycetes** | Tube-like, can form mycelia<sup>44</sup> | Peptidoglycan<sup>44</sup> | Heterotrophy<sup>44</sup> | Rejected, as *Prototaxites* had a clearly eukaryote-grade morphological and anatomical complexity<sup>45</sup>. |
| **Red, green or streptophyte algae** | Tube-like, septate with pores (red algae)<sup>46</sup> | Cellulose and various polysaccharides<sup>46</sup> | Photosynthesis exclusively<sup>46</sup> | Rejected, based on isotopic evidence that *Prototaxites* was heterotrophic<sup>13, 14</sup>, had a phenolic and aromatic composition distinct from that of algae (see text and<sup>16</sup>); and was terrestrial<sup>5,7</sup>. |
| **Land plants** | Typically box-like cells, develop distinct tissue types and diagnostic synapomorphies such as stomata, water and food conducting cells<sup>47</sup> | Cellulose, lignin, sporopollenin in spores | Photosynthesis | Rejected, as based on isotopic evidence that *Prototaxites* was heterotrophic<sup>13, 14</sup>, had a phenolic composition similar but distinct from that of land plants (see text), and no evidence for land plant synapomorphies<sup>48</sup>. |
| **Fungi** | Tube-like, septate with pores in some<sup>35</sup> | Chitin, chitosan and beta-glucan<sup>35</sup> | Saprotrophy<sup>35</sup> | Rejected, as *Prototaxites* is molecularly and anatomically distinct from fungi (see text), and no evidence of perylene (see text). |
| **Lichen** | Fungal symbiont is tube-like, can be septate with pores, presence of a photobiont<sup>35</sup> | Chitin, cellulose<sup>35</sup> | Mixotrophy<sup>35</sup> | Rejected, as *Prototaxites* is molecularly and anatomically distinct from fungi (see text), and no evidence of a photobiont<sup>7</sup>. |
| **Oomycetes** | Tube-like, can form mycelia<sup>48</sup> | Cellulose<sup>48</sup> | Heterotrophy<sup>48</sup> | Rejected, as phenolic composition of *Prototaxites* is distinct from that of oomycetes (see text), and major differences in morphology and complexity<sup>37</sup>. |
| **Metazoans** | Round cells without walls; typically organised into tissues | Chitinous cuticle (in some) | Heterotrophy | Rejected, as *Prototaxites* had a tube-like morphology, cell walls, and lacked metazoan synapomorphies such as distinct tissues<sup>49</sup>. |

## Acknowledgments

This work was supported by the Leverhulme Trust (ECF-2023-202; C.C.L.), the Royal Society (NIF\R1\211589; C.C.L., S.M. and RGS\R2\212063; A.J.H.), UK Research and Innovation Future Leaders Fellowship (MR/T018585/1; A.J.H.); Philip Leverhulme Prize (PLP-2023-324; A.J.H.); European Research Council (E.R.C.) under the European Union’s Horizon Europe research and innovation programme (grant agreement No 1101114969; S.J.); Science Foundation Ireland (22/PATH-S/10692; S.J.); Engineering and Physical Sciences Research Council grant (EP/Y037138/1; A.J.H.); and NERC E4 Doctoral Training Partnership (L.M.C., R.A.). We would like to thank Ivan Febbrari, Thin Sections and Lapidary Facility Manager, University of Edinburgh for the thin section preparation of *Prototaxites*; B. O’Connell (Cambridge University) for sedimentological advice; P. Orr (University College Dublin) for thin section preparation of Rhynie Chert; N. Fraser, A. Ross, and Y. Candela for assistance accessioning material into National Museums Scotland; F. Buckley and North Sea Core for their assistance with Rhynie chert specimens; and P. Kenrick and N. Clark for access to historic *P. taiti* collections. We thank Gianfelice Cinque and Diamond Light Source for access to the MIRIAM beamline, B22 (proposal number SM33471-1) that contributed to the results presented here. We thank M. Krings for useful discussion of *Prototaxites* and fossil fungi. This work was supported by funding for the Wellcome Discovery Research Platform for Hidden Cell Biology [226791] and we gratefully acknowledge support from the Microscopy core.

## Authors contribution

C.C.L., A.J.H., S.M. conceived the study and designed experiments. C.C.L. and A.J.H. coordinated the study. C.C.L., L.M.C., A.J.H., S.J., A.V.G., acquired the data with the help of S.M., L.P., H.V., R.A., E.R.D, A.T.B. Data were processed by C.C.L, L.M.C, and A.J.H. *Prototaxites* reconstruction were made by M.H. with inputs from A.J.H and C.C.L. C.C.L., A.J.H. and L.M.C. wrote the manuscript with inputs and reviews from all the other authors.

## Additional information

Supplementary information is available in the Supplementary Material and Edinburgh University DataShare.

## Data availability

Data for this work can be found in the manuscript, the supplementary material as well as on the Edinburgh University DataShare (doi: TBD). Fossil specimens described in this study are deposited in three collections: National Museums Scotland, UK, Lyon Collection, University of Aberdeen, UK, The Hunterian, University of Glasgow, UK.

## Methods

*Rhynie chert specimens*. NSC.36 was originally collected in farmland owned by the Windyfield Farm (NJ 349642 mE, 827852 mN) adjacent to the Site of Special Scientific Interest (SSSI) by a local landowner before 2021. Specimens were passed to North Sea Core to help distribute the samples for academic research with the mutual agreement of NatureScot. Blocks were distributed using the accession numbers North Sea Core NSC.01-NSC.45. NSC.36 was processed in the lab of AJH and the remaining sub-blocks of NSC.36 (**Fig. 1k**) are deposited in National Museums Scotland, UK (G.2024.5.1 and G.2024.5.2). Alongside our investigation of NSC.36 we also carried out a re-examination of the four thin sections that constitute the type material of *P. taiti* in The Hunterian, University of Glasgow (GLAHM Kid 2523 – 2526) as well as blocks of Rhynie chert containing *P. taiti* in the Lyon Collection, University of Aberdeen, UK (Lyon 156 and Lyon 48).

*Sample preparation.* NSC.36 was first cut in half (**Fig. 1j**) and a sub-block containing *P. taiti* was produced (**Fig. 1k**), from which six thin sections were made; three uncovered 100 µm thick specimens for FTIR (accessioned into National Museum Scotland (NMS) G.2024.5.3, G.2024.5.4 and G.2024.5.5) and three 30 µm thick thin sections with cover slips for light and confocal microscopy (initially named MPEG0056, MPEG0057 and MPEG0058, accessioned as NMS G.2024.5.6, G.2024.5.7, G.2024.5.8, respectively), the first two sections were taken from faces of the subblock perpendicular to each other to allow for the anatomy to be studied from different orientations. New covered and uncovered thin sections were made from blocks 156 and 48 containing *P. taiti* from the Lyon Collection, University of Aberdeen, UK. A total of four uncovered (UC) thin sections were created for FTIR analysis: Lyon 156 UC1, Lyon 156 UC2, Lyon 156 UCB1, and Lyon 48 UC1. A total of five thin sections with cover slips were made for light and confocal microscopy (Lyon 156 MPEG0071, Lyon 156 MPEG0072, Lyon 156 MPEG0073, Lyon 156 MPEG0078, Lyon 48 MPEG0070). Covered and uncovered thin sections of NSC36 were deposited in National Museums Scotland, UK. All new thin sections created from material from the Lyon collection were returned to the Lyon Collection, University of Aberdeen, UK.

*Photogrammetry.* For all three of the photogrammetry models (**Fig. 1d; Extended Data Fig. 2a** (Lyon 156), **Fig. 1i** (NSC.36 before cutting), and **1j** (NSC.36 after cutting)) a Canon EOS 5D Mark IV camera with a 100mm macro lens and tripod was used, with the following settings: ISO 100, f1/16, 1/5s. An adapted version of the photogrammetry protocol of Leménager et al.^50^ was used, in which the block was placed on an automated turntable (Genie Mini II and turntable) within a light box (Neewer) for even illumination. Each block had a series of 16 photos taken through a full rotation of the turntable, with the camera moved through four heights of the tripod (photos taken from ∼0°, ∼30°, ∼50° and ∼70° relative to the plane of the turntable) and a full rotation series of 16 photos taken at each of these heights. This full procedure (four heights, 16 photos at each height) was conducted twice for each block, resting first on one face then flipped to rest on the converse face (2.5 + 2.5 photogrammetry method). Scaled photogrammetry models were created using AgiSoft PhotoScan Professional and exported to CloudCompare to produce scaled renders.

*Optical microscopy.* Thin sections were studied and photographed using a Nikon ECLIPSE LV100D compound microscope (**Figs 1l**; **Figs 2a,e,h,I; Extended Data Fig. 2d**) and a Nikon SMZ18 Stereoscope (**Fig. 1e; Extended Data Fig. 2c**). **Fig 2e** was taken using Extended Depth of Focus (EDF Capture) in the NIS Elements software.

*Confocal Microscopy.* A Zeiss LSM 880 confocal microscope with Airyscan was used to produce regular and Airyscan CLSM images. Single plane Airyscan images (**Fig 2c,d; Extended Data Fig. 2e,f**) were taking using a x40 oil immersion lens, 488nm and 561nm lasers, long pass emission window, 2.5Au pinhole, pixel size 0.05 µm, 2576×2576 frame size. Regular CLSM settings (with all other parameters the same as those used for Airyscan with the exception of a 1 Au pinhole) with image stitching was used to create the image in **Fig 2b**. Z-stack images were produced with all settings as for single plane Airyscan CLSM imaging, with a 0.27 µm interval between successive images, corresponding to a total of 114 slices.

*3D reconstruction.* The Airyscan CLSM z-stack of the medullary spot region was first opened in Fiji^51^, subjected to image contrast enhanced on all slices, colour inverted, then saved as a jpeg file series. 3D reconctruction was conducted using SPIERS^52^. The jpeg file series was imported to SPIERSedit, and a single large axis was picked to reconstruct branching (axis in top right of **Fig 2f**). After application of automatic thresholding to the image series, the branching pattern was traced through the image series and segmented using curves, the thresholded image was then adjusted manually, and masks were produced from curves (illustrated in blue on **Figs. 2f, g**). Images were rendered in Blender, with adjustments to light, shading and colour for clarity of the resulting render (**Fig 2g**).

*Transmission benchtop FTIR.* Small fragments of the NSC.36 block containing only *Prototaxites* were macerated in acid to remove silica following a low-manipulation protocol adapted from^53^, in the Oceanography Clean Laboratory (School of Geosciences, University of Edinburgh). The samples were placed in an 120ml Teflon jar and bathed overnight in hydrochloric acid (HCl) to remove potential carbonate influence (no reaction was noted). HCl was discarded, and the samples were left for a week in analytical grade hydrofluoric acid (HF) with daily light hand-agitation of the jar. After complete dissolution and sedimentation, the acid surplus was removed, and samples were brought to boil with HCl for half a day to remove or prevent the formation of secondary minerals. The acid was then neutralized by successive washes with water.

A drop of macerate was deposited on a zinc selenide window, placed to dry in an oven at 60°C for 2H, and a spectrum was acquired in the range 4000-650 cm^-1^, with a spectral resolution of 4 cm^-1^ and 16 accumulations, using an AlphaII FTIR spectrometer (Bruker) at the UK Centre for Astrobiology (School of Physics and Astronomy, University of Edinburgh). Spectra were pre-process using Quasar 1.9.2 (Orange^54,55^) (atmosphere-corrected, Gaussian smoothing and baseline correction). A second derivative spectrum was obtained by applying a Savitzky-Golay filter of order 2 and window 21. Band assignments are shown in **Supplementary Table 1**.

*Attenuated Total Reflectance-FTIR (ATR-FTIR).* ATR-FTIR was conducted at room temperature on a Smiths IlluminateIR microscope equipped with a liquid nitrogen-cooled MCT detector and a diamond coated ATR objective (magnification x36) at the School of Chemistry (University of Edinburgh). Backgrounds were taken in air before analysis. We acquired the spectra in reflection mode by combining 128 accumulations in the range 4000–650 cm^−1^ at a resolution of 4cm^-1^ with the software Qual ID 2.51 (Smiths). We processed the spectra using software Quasar. We truncated the spectra to analyse the range between ca. 1445-1750 cm^-1^ and 2760-3000 cm^-1^, removing intervals due to hydroxyl absorptions (4000–3000 cm^−1^), intense vibration of silica (1400–650 cm^−1^), atmospheric CO_2_, ATR diamond absorption and high wavenumber silica overtones (2760-1750 cm^-1^). We corrected the baseline, applied a light Gaussian smoothing. Finally, to minimize the influence of thickness and differences in concentration of organic matter, the spectra were normalised on the highest silica overtone absorption peak (1615 cm^-1^). For inspection and band assignment, average second derivative spectra for each domain (chitinous and non-chitinous) were obtained by applying a Savitzky– Golay filter (**Supplementary** Figs. 1,2). Band assignements are shown in **Supplementary Table 1**. Additional spectra of Rhynie chert samples of plants, fungi, animals, peronosporomycetes and amoeba were taken from^31^.

*Data exploration – features selection*. After preprocessing, the spectra were organised into four datasets: (1) Prototaxites vs Bacteria; (2) Prototaxites vs Fungi; (3) Prototaxites vs Chitinous organisms (Fungi + Arthropods); and (4) Prototaxites vs Plants (including plant spores). A fifth dataset comprising the 102 samples from Plants (n=37), Fungi (n=24), Arthropods (n=12), Bacteria (cyanobacteria; n=10), Peronosporomycetes (n=4), Amoebae (n=3), and *Prototaxites* (n=12) is also compiled for one-class classification (see below). For datasets 1-4, each sample was one-hot encoded according to their biological group. These formatted datasets were then processed through the pipeline illustrated in **Extended Data Fig 3**. Analyses were carried out in Python 3.12.8. Data exploration began by exploring the data structure with Principal Component Analysis (PCA). PCA is a fundamental step before performing supervised classification tasks because it performs a reliable dimension-reduction that allows selection of robust features for classification whilst remaining easily interpretable. PCA was conducted using the scikit-learn package. We extracted the first ten Principal Components (PCs) with their respective Explained Variance and Cumulative Variance. To control the robustness of our PCA, we compiled scree plots, scores and loading spectra, and tested the stability of each PCA by calculating the cosine angle between the original dataset and resampled dataset using bootstrapping (100 bootstraps). A cosine angle close to 1 for a PC indicates low variation and shows that the PCA is not sensitive to variation in the dataset for this PC (i.e., this PC represents a real pattern in the data)^56^. Finally, using the most stables PCs as references we performed outlier detection using Hotelling’s T^2^ versus Q residual values^57,58^. Detected outliers were removed and each dataset was recompiled. After these first robustness checks, we performed new PCA analyses on the recompiled datasets. Based on the robustness checks for these new analyses, we selected PCs for further supervised learning that retained sufficient variance and represented interpretable biological information (see **Extended Methods**). The retained PCs (PC1 and PC2 for dataset 1 and PC1-PC4 for datasets 2-4) were directly used as variable for classification tasks (see below). Finally, we extracted the intensities of the main biologically informative absorption bands in the spectra, based on the loading spectra of the retained PCs. These bands (bands 1, 2, 7 and 10-12 in **Supplementary Data Table 1**) were used for correlation analysis with Canonical Correspondence Analysis (CCA) and for further classification tasks as a comparison with the full spectra approach described above.

*Data Exploration – CCA.* CCA is a multivariate supervised statistical method used to explore the correlation between a matrix of response variables and a set of explanatory variables^59,60^. CCA was conducted on the extracted bands using the software PAST 4.17. We used our lineages as response variables (with each sample one-hot encoded) and the extracted band intensities as explanatory variables. Results of the ordination and correlation between the lineages and the spectral feature can be seen in **Fig 3**.

*Modelling.* Multiclass supervised classification tasks were performed according to the pipeline presented in **Extended Data Fig 3.** Modelling was performed in Python 3.12.8 using the scikit-learn package on the datasets without outlier. To keep the models simple, we performed binary classifications. Similarly, we follow a parsimony principle by testing first simple linear models (Linear Discriminant Analysis, LDA) before moving to more complex algorithm (Support Vector Machine, SVM)^58^. Data were randomly split into training (70%) and test set (30%), whilst retaining the ratio of each lineage (stratified). After splitting, the training data were subjected to dimension reduction using PCA retaining a number of PCs determined by data exploration steps. The training dataset for datasets 2-4 show a strong data imbalance between each class. To address this imbalance, we performed Synthetic Minority Oversampling Technique (SMOTE), which generates synthetic examples for the minority class^61^. PCA and SMOTE are performed after data splitting to avoid data leakage, which would compromise the test set and the robustness of the models. For all analyses, validation was perform using Leave-One-Out-Cross-Validation on the training set, a methods well adapted for small dataset^62, 58^. After cross-validation, the model was run on the whole training set. For each analysis, learning curves were computed to control if the size of our datasets were sufficient to obtain robust results. For each LDA, we computed score plots and controlled robustness using bootstrap stability tests, confusion matrix and five performance metrics (Accuracy, Precision, Recall, F1 and Matthew Correlation Coefficient (MCC)). Each SVM analysis followed the same robustness checks. For SVM we also computed the two-dimensional decision boundary based on the best parameters (C and gamma) obtained after grid-search during cross-validation.

LDA performed well for dataset 1, scoring 1 for each parameter in both the training and the test set. Other datasets showed improved performance and robustness with SVM, consistently scoring between 0.91 and 0.96 for each of the metrics and above 0.8 for MCC in both training and test sets (see details in **Extended Methods**). Unlike full spectra, models using selected individual bands performed poorly, showing a strong discrepancy between training and test set performances, especially lower MCC.

Finally, one-class modelling was used to show that *Prototaxites* is not only molecularly different from the Rhynie chert bacteria, plants, and chitinous organisms when treated separately, but also from all Rhynie chert fossils taken together (dataset 5). We performed one-class modelling using Data Driven Class Analogy (DD-SIMCA) as developed by Kucheryavskiy and colleagues^63^ (www.mda.tools/ddsimca). We trained a PCA model on the *Prototaxites* samples (n=12). To mitigate the effect of the small sample size, we use resampling Leave-One-Out-Cross-Validation. We used a rigorous feature selection approach based on sensitivity, and less permissive outlier detection using robust parameter estimates on the training set^63,64^. No outliers were detected. We tested all non-*Prototaxites* fossils with the model trained on *Prototaxites* samples. The model provided an excellent true negative rate (specificity) of 0.911, indicating high confidence that *Prototaxites* differs from all other Rhynie chert fossils in its fossilisation products. *Synchrotron FTIR.* A double polished thin-section (80 μm) was produced from *Prototaxites* NSC.36 (School of Geosciences, University of Edinburgh) and analysed at the B22 beamline of Diamond Synchrotron (UK). Two maps were acquired through a medullary spot and the body, and one line was acquired through a type 2 tube. Using Quasar, spectra were atmosphere corrected and truncated between 3000 and 1440 cm^-1^. Second derivatives were calculated using a Savitzky-Golay Filter of order 2 and window 9. We applied a light Gaussian smoothing and a vector normalisation.

975 individual spectra were extracted from the maps and line, and labelled as: body (i.e. dense pack of type 1 and 2 tubes) and medullary spot (mix of dense tiny tube and branching larger tubes). We did not found informative difference between the 2 groups (see **Supplementary** Fig. 5). The pre-processing of the spectra was repeated using the steps above, but with baseline correction instead of calculation of the second derivative. CH_2_ asymmetric stretching bands at ca. 2925 cm^-1^ were integrated together with CH_3_ asymmetric stretching bands at ca. 2960 cm^-1^, and a CH_3_/CH_2_ semi-quantitative ratio was calculated based on band intensity. This ratio provides information about carbon chain length and branching; the lower the ratio, the higher the chain-length and the more limited the branching (based on the construction of alkane chain with CH_3_ at the extremity and CH_2_ forming the chain, e.g.,^31,34,65–67^). We calculated average ratio values of 0.75 for the body and 0.71 for the medullary spots, which both correspond to chain length of ca. nine carbons^67^.

*Biomarkers.* For biomarker analysis, samples of block NSC.36 were separated into those consisting of bulk material absent *P. taiti* and those consisting entirely of *P. taiti,* to enable comparative analysis. A standard clay brick was used as a control material (furnaced at 550 °C for 12 hrs and underwent every step of the sample preparation, extraction, and analysis that the NSC.36 samples were subjected to). All glassware, ceramic ware, and aluminium foil was furnaced at 550 °C for 12 hrs. Procedural controls containing no material were run alongside every NSC.36 and combusted brick sample from extraction to analysis, in order to control for contamination introduced in the laboratory. Samples were prepared for extraction in the Earth Surface Research Laboratory at Trinity College Dublin. Large pieces were first broken down to smaller pieces in a jaw crusher and subsequently milled to a powder using a ball mill. Components of the jaw crusher and milling equipment that came into contact with the sample material were cleaned with deionised water, methanol, and chloroform between uses. Smaller fragments of *P. taiti* were crushed to a powder using a mortar and pestle.

Samples were weighed into 50 mL glass centrifuge tubes with PTFE lids (ca. 13 g for bulk and brick, ca. 3 g for Prototaxites). 100 μL 100 ppm squalane was added to the combusted brick and procedural control in order to estimate % recovery. Samples and controls were ultrasonically extracted three times with 20 mL 9:1 chloroform:methanol at 40 °C for 15 minutes. Following each extraction, tubes were centrifuged at 3000 rpm for 30 minutes. Extracts were transferred to round bottom flasks with a glass Pasteur pipette. Combined extracts were then filtered through glass fibre filters (Whatman GF/A) before being concentrated under anhydrous N_2_. Concentrated extracts were transferred to 2 mL GC vials, dried under anhydrous N_2_, and subsequently resuspended in 100 μL extraction solvent. 100 μL 100 ppm 5α-cholestane was added to the combusted brick and procedural control as an internal standard.

Samples were extracted in triplicate by ultrasonication with 9:1 (v/v) chloroform:methanol at 40 °C. Total lipid extracts were combined and concentrated under anhydrous N_2_ prior to analysis. Aliquots (1 µl) of samples were injected in triplicate onto an Agilent model 7890N gas chromatograph coupled to an Agilent 5973N mass selective detector operating in electron impact mode at 70 eV. The column was a 30 m HP-5MS column (0.25 mm i.d., 1 μm film thickness). Each sample was injected with a 2:1 split ratio. The GC inlet temperature was 280 °C and the oven programme was 65 °C (held 2 min) to 300 °C (held 20 min) at 6°C/min. Each sample was run in full scan and selected ion monitoring (SIM) modes (m/z 252 for perylene). Individual compounds were assigned for comparison with mass spectral library databases (NIST98 and Wiley275) and comparison of mass spectrometry patterns with published spectra.

---

**Extended Data Fig. 1.**
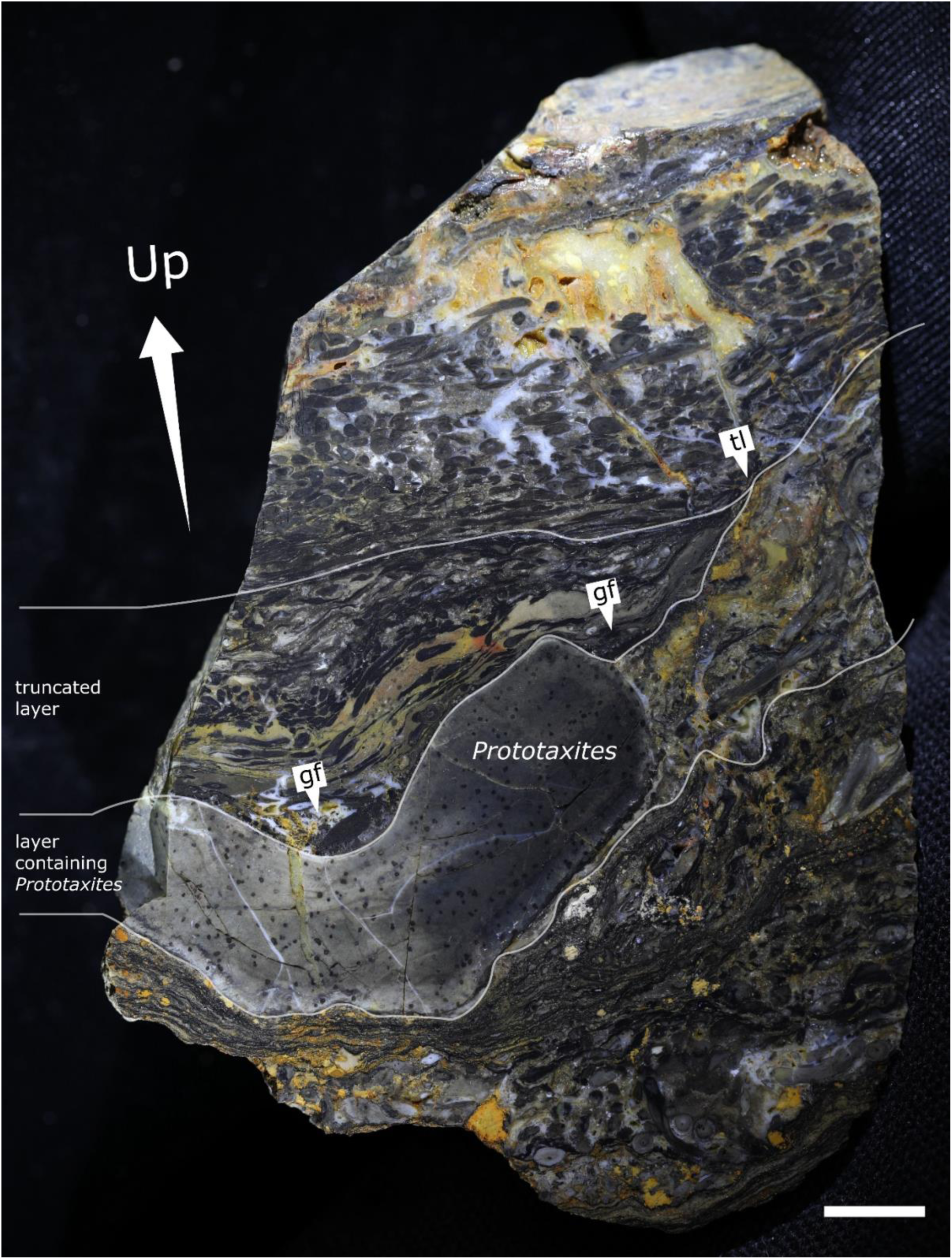
| Establishing the orientation of NSC. 36. Annotated view of Rhynie chert block NSC.36 after being cut in half to show the location of the specimen relative to surrounding sediment, and interpret its orientation. Selected laminations and geopetal features, which support the way-up direction indicated: tl = truncated lamina; gf = geopetal sediment fill. Scale bar: 5 mm.

**Extended Data Fig. 2.**
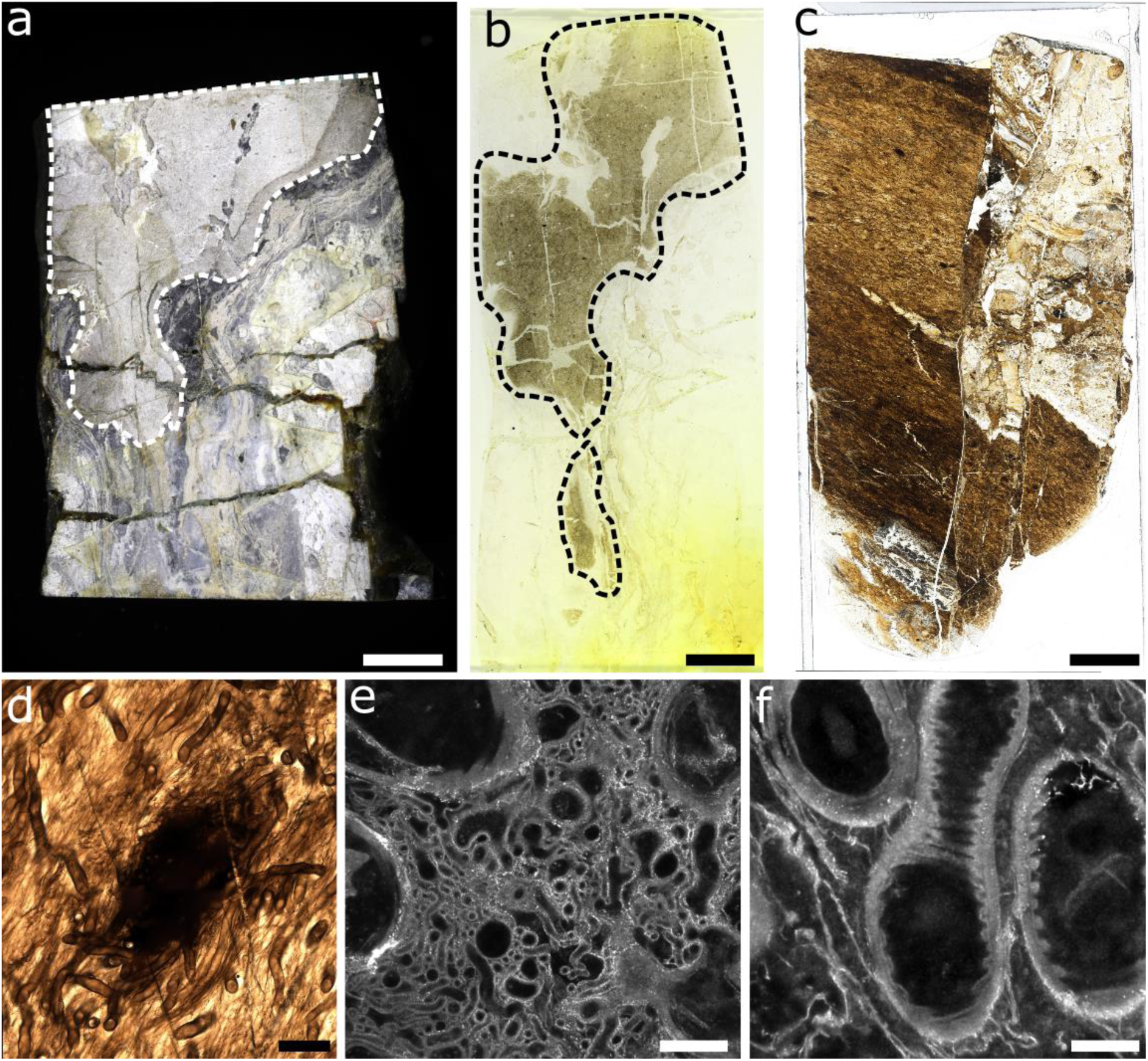
| Anatomical features of *Prototaxites taiti* from Lyon Block 156. **a-c**, Lyon 156 block (**a**) and peel (**b**) with *P. taiti* highlighted in dashed lines. Slide produced from the block in (**a**), showing darker tissue than the NSC.36 specimen (**c**). **d-f**, Anatomical investigation of Lyon 156 *P. taiti*, showing a spot region in light microscopy (**d**), Airyscan CLSM imaging of the spot region in (**d**) (**e**), Airyscan CLSM imaging of tubes with annular thickenings in the body of Lyon 156 (**f**), these tubes are notably thicker than those seen in NSC.36 (and the type material) but similar to the thickened tubes seen in other *P. taiti* species. Scale bars: 1cm (**a,b**), 500µm (**c**), 200µm (**d,e**), 20µm (**f**). Specimen accession codes: Lyon 156 (**a**), Lyon Peel 156/1 (**b**), Lyon 156 MPEG0078 (**c-f**).

**Extended Data Fig. 3.**
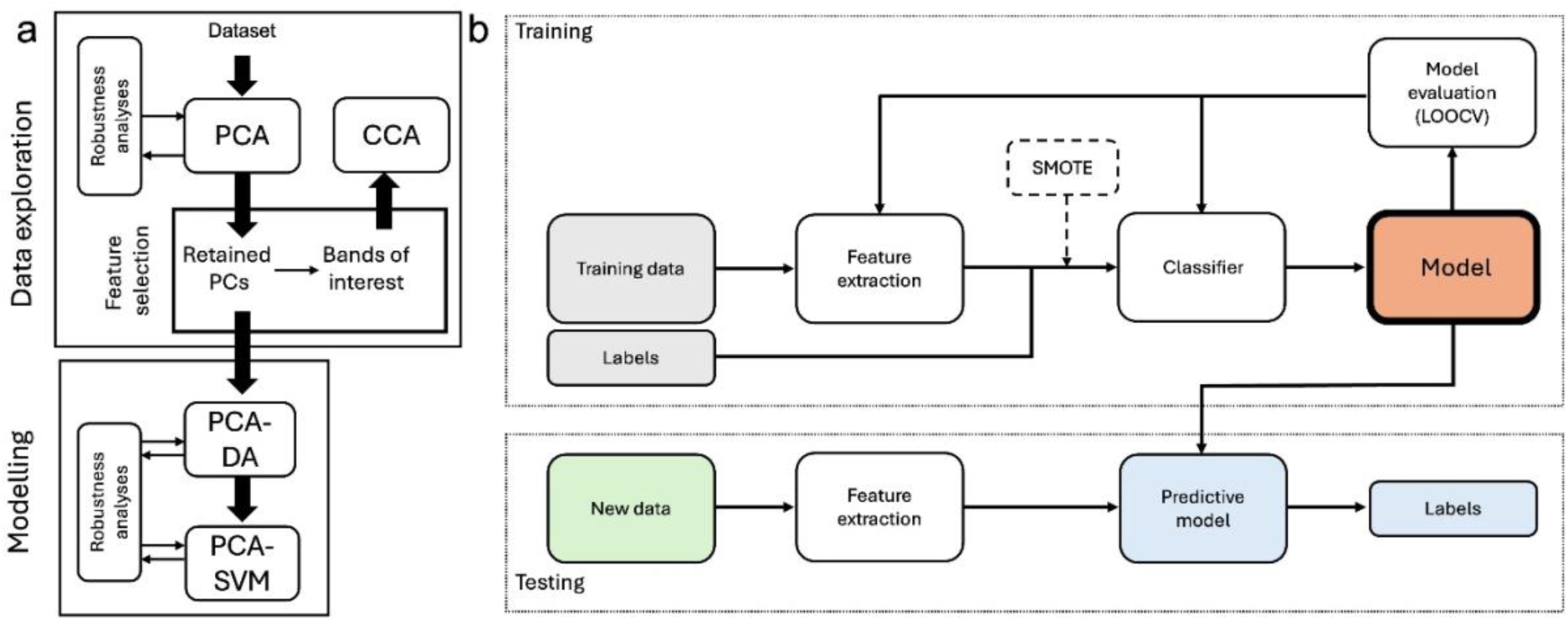
| Analytical pipeline used in this study. **(a)** illustrates the key steps of our workflow, comprising data exploration (dimension reduction and feature selection using Principal Component Analysis (PCA), and Canonical Correspondence Analysis (CCA) for correlation and ordination) and modelling using Linear Discriminant Analysis combined with PCA (PCA-DA) and Support Vector Machine combined with PCA (PCA-SVM). **(b)** details the modelling step with the splitting of the data for training and testing. Training consists of feature extraction (the PCA stage) followed the correction for class-imbalance (SMOTE) if needed and the choice of classifier (DA or SVM). The model obtained is cross-validated by Leave-One Out-Cross-Validation (LOOCV). Testing consists of a feature extraction stage following the same parameters than the training stage and a prediction stage where the label (*Prototaxites* or plant, fungi, bacteria) for new, unseen instances in the test set are predicted by the cross-validated model.

**Extended Data Fig. 4.**
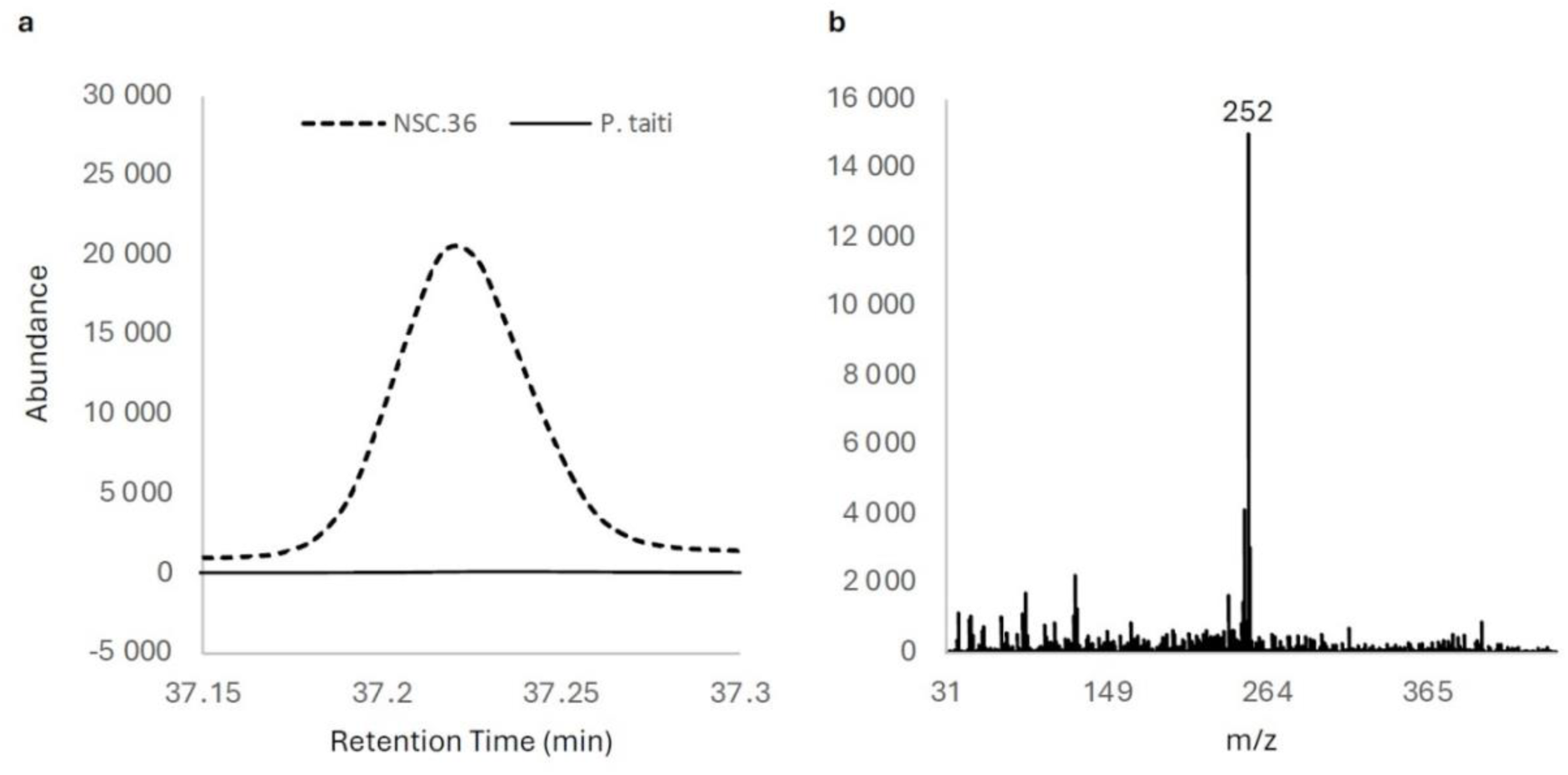
| Perylene was not identified in *Prototaxites taiti.* Perylene detected in bulk NSC.36 material but not in pure *P. taiti*. **a**, Chromatogram trace showing possible ascomycete biomarker perylene peak with a retention time of ca. 37.22 minutes for the bulk NSC.36 material (dashed line) and the pure *P. taiti* material (solid line). Chromatogram was recorded in selected ion monitoring mode (SIM) at m/z 252 which is diagnostic for perylene. Only the relevant chromatographic region is shown. B, mass spectrum of perylene showing diagnostic m/z 252 ion.

